# Bilayered Biofabrication Unlocks the Potential of Skeletal Muscle for Biohybrid Soft Robots

**DOI:** 10.1101/2024.03.01.582738

**Authors:** Aiste Balciunaite, Oncay Yasa, Miriam Filippi, Mike Y. Michelis, Robert K. Katzschmann

**Affiliations:** Soft Robotics Lab, Department of Mechanical and Process Engineering, ETH Zurich

## Abstract

The emerging field of biohybrid robotics aims to create the next generation of soft and sustainable robots by using engineered biological muscle tissues integrated with soft materials as artificial muscles (bio-actuators). Both cardiac and skeletal muscle cells can be used for biohybrid actuation. Generally, cardiac bio-actuators take the shape of thin cellular films, while locomotive skeletal muscle bio-actuators form bulk tissues. The geometry of a bio-actuator should be optimized for the type of desired motion, *e.g*., thin film layers are optimal for swimming actuators mimicking fish. Until now, the geometry of skeletal muscle bio-actuators has been constrained to ring- or block-like tissues generally differentiated around a pair of pillars due to the need to oppose the contraction force exerted during the skeletal muscle differentiation process. In this work, we extend the possible geometry of skeletal muscle bio-actuators by demonstrating a bilayered design that mimics the motion of jellyfish. We take advantage of a volumetric printing method, *i.e*., xolography, which allows us to micropattern poly(ethylene glycol) diacrylate and gelatin methacrylate hydrogels to serve as scaffolds for seeding a layer of the skeletal muscle cell matrix. We demonstrate the locomotion speed of our bio-actuators is 3.4x faster than previously reported counterparts. In addition, our skeletal bio-actuators outperform most cardiac ones. Further optimization of our bilayer biofabrication for improved reproducibility of the maturation process of the skeletal muscle tissue will pave the way for the next generation of performant skeletal muscle-based actuators for biohybrid robots.

## I. Introduction

Biohybrid soft robotics is an emerging active research field that exploits the natural abilities of muscle cells integrated with artificial components for developing the next generation of soft robots [1], [2]. Biohybrid soft robots promise advanced functionalities, such as high energy conversion efficiency and self-healing, which are challenging to capture in traditional soft robots [3]. Biohybrid soft robots rely on nature’s actuator, *i.e*., muscle cells, to realize motion. While either cardiac or skeletal muscle cells can be used as natural materials for bio-actuation, skeletal muscle cells have the advantage of being more controllable in that their biomechanical response depends on applied stimuli. Moreover, these cells are available as cell lines, easily expanded, and suitable for high-throughput biofabrication of larger bio-actuators.

Despite the potential of skeletal muscle cells, the most performant biohybrid swimmers have still been realized from cardiac bio-actuators with bioinspired planar geometries created from either a single cell layer (*i.e*., termed as *thin films* [4]) or hydrogel-based cell layers. These geometries enable the robots to efficiently move within fluids by mimicking the swimming motions of aquatic animals [5]–[7].

A key reason that cardiac cells have been used as effective bio-actuators is that cardiac bio-actuators do not need to follow a differentiation process after fabrication to achieve contraction. In contrast, engineered skeletal muscle tissue needs to follow a 14 day differentiation process to fuse into multinucleated, contractile myofibers. To this purpose, skeletal muscle bio-actuators need to grow in the presence of topographical stimuli and skeleton-like structures that provide the necessary mechanical cues and can withstand the shrinking forces of the forming skeletal muscle tissues [8]– [11]. The thin film scaffolds used with cardiac cells are not robust enough to withstand the full skeletal muscle differentiation process without losing structural integrity. Hence, skeletal muscle bio-actuators are mostly realized from 3D bulky tissue geometries stretched over pillar-like structural components. These bulk tissue designs are not optimized for motion in liquid environments, thus limiting their ability to locomote efficiently. Despite the potential of skeletal muscle, no biohybrid soft robot with a planar design for efficient and biomimetic locomotion has ever been realized from skeletal muscle cells.

## II. Bio-Actuator Design and Fabrication

We aimed to adapt performant cardiac biohybrid locomotion geometries to use with skeletal muscle cells. We designed a bilayered planar actuator inspired by the locomotion of previously reported jellyfish or ray-like bio-actuator morphologies [5], [6]. The curved bio-actuator moves by contracting inwards, similar to a scallop motion [12]. Unlike previously reported cardiac planar bio-actuators, we aimed to take advantage of the controllability and tissue formation of skeletal muscle compared to cardiac muscle. We developed a rapid, easily customizable method to grow skeletal muscle tissue molded onto a pre-fabricated scaffold. We measured the mechanical properties of the scaffolds to inform simulation experiments of scaffold bending. Finally, we characterized the swimming behavior of our bio-actuators (Figure 1).

**Fig. 1.**
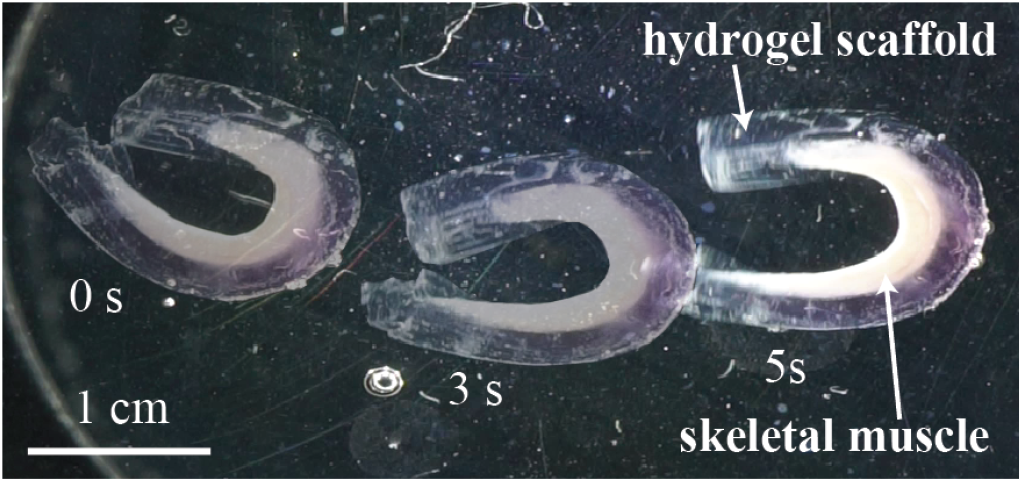
Bilayered bio-actuators enable the use of cultured skeletal muscles to achieve fast swimming motions in centimeter-scaled soft robots.

**Fig. 2.**
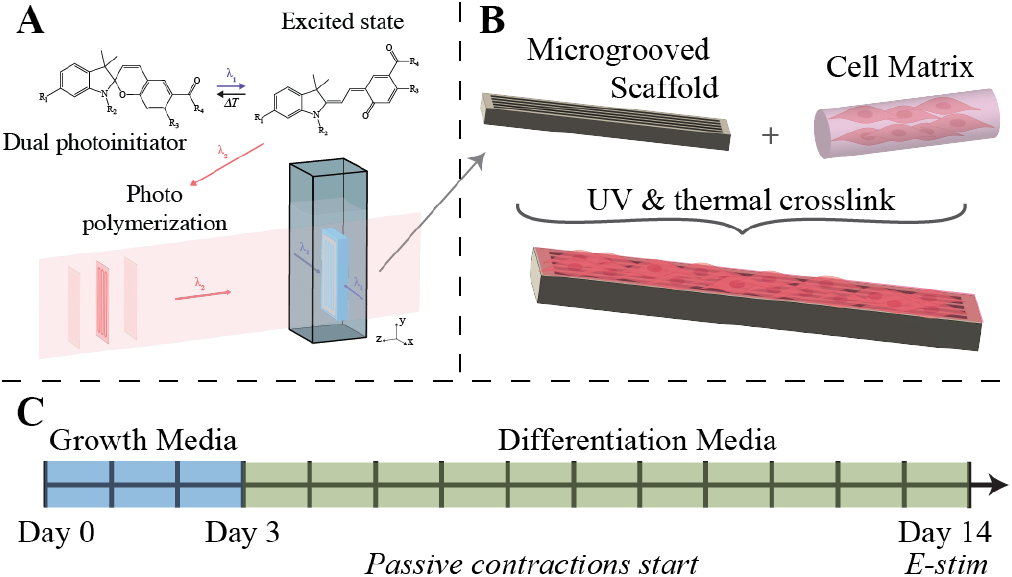
Overview of the bilayered skeletal muscle bio-actuator fabrication. (A) 3D printing of micro-grooved hydrogel scaffolds with xolography. (B) Integration of skeletal muscle cells with the 3D-printed scaffolds for creating the bio-actuators. (C) Differentiation and maturation timeline of skeletal muscle bio-actuators.

To facilitate macroscopic movement, the scaffold needs to be soft and contain cell adhesion cues. Therefore, we used a hydrogel composed of poly(ethylene glycol) diacrylate with a M.W. of 700 (PEGDA700) and gelatin methacrylate (GelMA). PEGDA700 provides patternability and structural stability during the differentiation process to the hydrogel. GelMA adds softness and contains cell adhesion sites. To realize microscale hydrogel topography in a fast and scalable way, we used xolographic volumetric printing [13].

Figure 7 shows the general fabrication scheme of the bio-actuators. First, the scaffolds (1 mm x 3.5 mm x 15 mm, grooves: 300 μm width x 500 μm depth x 1.4 mm) are fabricated using a xube volumetric printer. The muscle cell matrix layer is then cast onto the scaffold, and the bio-actuator is matured for a period of 14 days at 37 °C and 5% CO_2_. After following the maturation protocol, the bio-actuators are tested for contractility and locomotion characteristics by application of a pulsed direct current electric field (3.3 V/cm, 1% duty cycle, 1 - 5 Hz).

### Xolographic printing of scaffolds

The hydrogel scaffolds were fabricated using a xube volumetric printer (xolo3D GmbH, Berlin, Germany). The hydrogel material was mixed from 15 vol% PEGDA700 (Sigma-Aldrich, St. Louis, MO), 5 wt% GelMA (prepared in-house), and 0.25 wt% photoinitiator (xolo3D GmbH, Berlin, Germany) dissolved in 1M Bis-Tris (Sigma-Aldrich, St. Louis, MO). After printing, hydrogels were post-processed by soaking in phosphate-buffered saline (PBS) for 30 min to remove non-polymerized polymer residues from the channels, followed by a UV curing at 365 nm for 120 s to ensure complete polymerization. Scaffolds were soaked in PBS overnight to equilibrate to pH 7.4 and sterilized under UV for 15 min before use.

### Muscle Layer Fabrication

C2C12 cells (mouse myoblasts, American Type Culture Collection) were cultured in a growth medium (GM) composed of high-glucose Dulbecco’s modified Eagle’s medium (DMEM) supplemented with 10% (v/v) heat-inactivated fetal bovine serum (FBS) and 1% (v/v) penicillin-streptomycin (PS) until they reached approximately 80% confluency, then passaged. Cell matrix was prepared by suspending 10^6^ cells/mL in 30% Matrigel (Corning, Lot #3068003), 1.6 mg/mL collagen (Cosmo Bio Co., Tokyo, Japan), and 5% GelMA. A volume of 20 μL of cell-matrix solution was deposited onto prepared scaffolds, then the constructs were UV polymerized at 405 nm for 30 s. Constructs were incubated for 30 min to allow complete gelation of Matrigel and collagen before adding GM supplemented with 1 mg/mL 6-Aminocaproic acid (ACA) to modulate degradation and remodeling of the hydrogel by cells. After 3 days of growth, constructs were switched to differentiation medium (DMEM, 10% (v/v) horse serum, 1% (v/v) PS, 1 mg/mL ACA, 50 ng/mL insulin-like growth factor 1 (IGF-1)) and cultured for 10 more days.

### Tissue Staining

After *in vitro* culture, the bio-actuators were fixed for 1 h in a 4% (v/v) paraformaldehyde solution at room temperature. Then, they were washed in PBS (5 min, 3x) and stained for DAPI, F-actin, and Myosin Heavy Chain (MyHC). Briefly, samples were permeabilized with 0.1% Triton X-100 for 20 min and blocked with a 1% bovine serum albumin solution for 1 h. Bio-actuators were stained with 4’-6-diamidino-2-phenylindol (DAPI, dilution 1:1000; 10 min); AlexaFluor 488 phalloidin (#R37110, Invitrogen) for 1 h; and mouse anti-MyHC antibody (Myosin 4, eFluor 660, Clone: MF20, Affymetrix eBioscience) for 4 h. All images were taken using a confocal microscope (Nikon Ti2).

## III. Experimental Results

### A. Modeling Scaffold Geometry to Optimize Deformation and Locomotion

Due to the quick and customizable nature of xolographic printing, we could fabricate scaffolds of various size ranges with the same grooved topography. We aimed to achieve a macroscopic deformation of the scaffold upon contraction of the muscle layer that would be sufficient to result in locomotion. To inform our experimental designs, we constructed a simplified finite elements model of the layered planar bio-actuators using the structural mechanics module of COMSOL Multiphysics 6.1 (Figure 3). The scaffold was modeled as a linear elastic material with a variable Young’s Modulus and Poisson’s ratio of 0.49. The muscle material was taken to have a Young’s Modulus of 25 kPa and a Poisson’s ratio of 0.49 based on literature [14]. The bio-actuator was modeled using half of the geometry with the center fixed. The muscle layer was taken to be 300 μm in thickness and assumed to span the length of the hydrogel scaffold. The scaffold-muscle adhesion was captured by using a union. The force of the muscle produced during the contraction was approximated as a unidirectional body force parallel to the grooves of the scaffold.

**Fig. 3.**
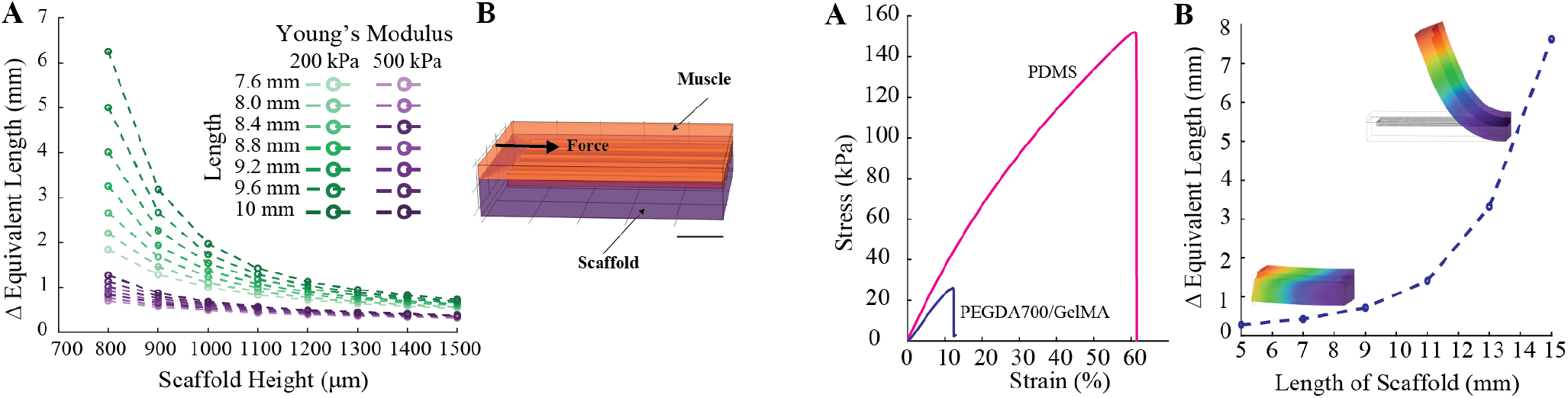
FEM modeling of a layered planar actuator. (A) Effect of changing scaffold thickness and length on bio-actuator deformation. Bio-actuator deformation is shown as the change in equivalent (horizontal) length of the fully bent bio-actuator. (B) Symmetric bending is modeled by using one-half of the bio-actuator geometry and defining a scaffold (purple) and muscle (orange) material. Unidirectional force is applied as a body load on the muscle in the direction of contraction. Scale bar is 1 mm.

Figure 3 shows that increasing Young’s Modulus of the scaffold, increasing scaffold thickness, or decreasing scaffold length results in a small macroscopic deformation of the scaffold upon contraction with the same force. Crucial considerations from a design perspective were that decreasing scaffold thickness ≤ 1 mm and optimizing the scaffold material stiffness with scaffold length would have a large effect on the deformation amount.

To experimentally investigate the effects of scaffold size, we manufactured scaffolds with 1 mm thickness, width of 3.5 mm, and lengths of 5 mm and 15 mm. The hydrogels swelled 140% after exposure to water, reaching final dimensions of 1.4 mm thickness, width of 5 mm, and lengths of 3.6 mm and 21 mm, with grooves of 420 μm. We determined the Young’s Modulus of our PEGDA700/GelMA scaffold to be 200 kPa based on tensile testing (Figure 4A). Based on COMSOL simulations, we expected that bio-actuators of 5 mm length would deform only 0.5 mm, while bio-actuators of 15 mm length would have deformation of approximately 7.5 mm (Figure 4B). Experimentally, we characterized that the 5 mm bio-actuators changed their equivalent length by 0.3 mm, while 15 mm bio-actuators changed their equivalent length by 4.5 mm. The observed discrepancy at 15 mm could be partially explained by the deformation of the real 15 mm bio-actuators starting from an already bent position (unlike in the simulation).

**Fig. 4.**
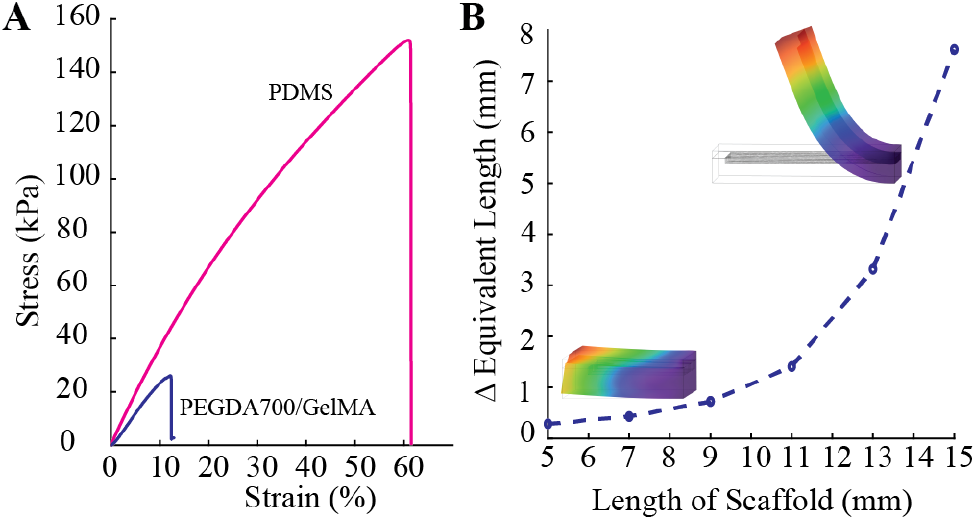
Scaffold material and size considerations. (A) Tensile testing results of our PEGDA700/GelMA hydrogel material and 1:20 PDMS (previously used for cardiac swimmers [4]–[6]). (B) COMSOL simulation presents the relation of increasing the length of the scaffold while keeping E = 200 kPa and thickness = 1 mm as manufactured.

### B. Locomotion Behavior of Bio-Actuators

To achieve contractility, bio-actuators were fully submersed in approximately 1 cm of liquid and electrically stimulated with direct current pulses (E = 3.3 V/cm, 1% duty, 1 - 5 Hz) using a function generator (DG812, Rigol, Beijing, China). Videos were captured from above at 25 fps using a darkfield illumination and a Sony α6000 camera with a macro lens (Sony SEL50M28). The motion of the bio-actuators was analyzed stroke-by-stroke using FIJI [15]. Average fabrication batch sizes were 5-7 actuators. However, reproducibility of swimming speed was a significant challenge that will be discussed in the context of optimizing the maturation protocol. A key point of the maturation protocol was the matching between the passive contractile force of the differentiating skeletal muscle tissue with the opposing force to bending provided by the hydrogel scaffold. Due to variability in the muscle tissue differentiation, not all scaffolds achieved the optimal bending amount to achieve efficient propulsion through liquid. All scaffold started in a flat configuration, and changed shape over the differentiation cycle. Some would curl onto themselves, while others would remain flat. The following investigation into bio-actuator swimming characteristics is shown for the fastest bio-actuator, which achieved efficient motion by curving into a c-shape.

Figure 5 shows the distance traveled by an optimal bio-actuator (Figure 5A) at different electrical stimulation frequencies, with the corresponding velocity values shown in Table I. Increasing the actuation frequency increased speed. Though the absolute speeds at 4 and 5 Hz were different, the distance over time graph in Figure 5B shows that the slope of both the 4 and 5 Hz lines is similar and that the two paths diverge only at the end. Congruently, the distance traveled per stroke of the bio-actuator at 4 and 5 Hz is similar (Table I).

**TABLE I.**
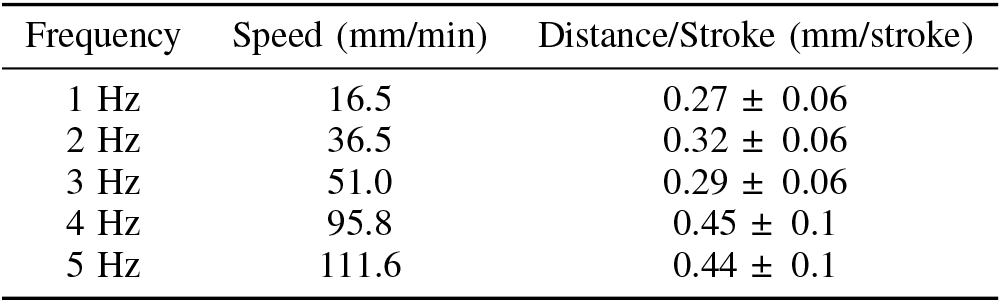
Characteristics of the bio-actuator at varying stimulation frequencies.

| Frequency | Speed (mm/min) | Distance/Stroke (mm/stroke) |
| --- | --- | --- |
| 1 Hz | 16.5 | $0.27 \pm 0.06$ |
| 2 Hz | 36.5 | $0.32 \pm 0.06$ |
| 3 Hz | 51.0 | $0.29 \pm 0.06$ |
| 4 Hz | 95.8 | $0.45 \pm 0.1$ |
| 5 Hz | 111.6 | $0.44 \pm 0.1$ |

**Fig. 5.**
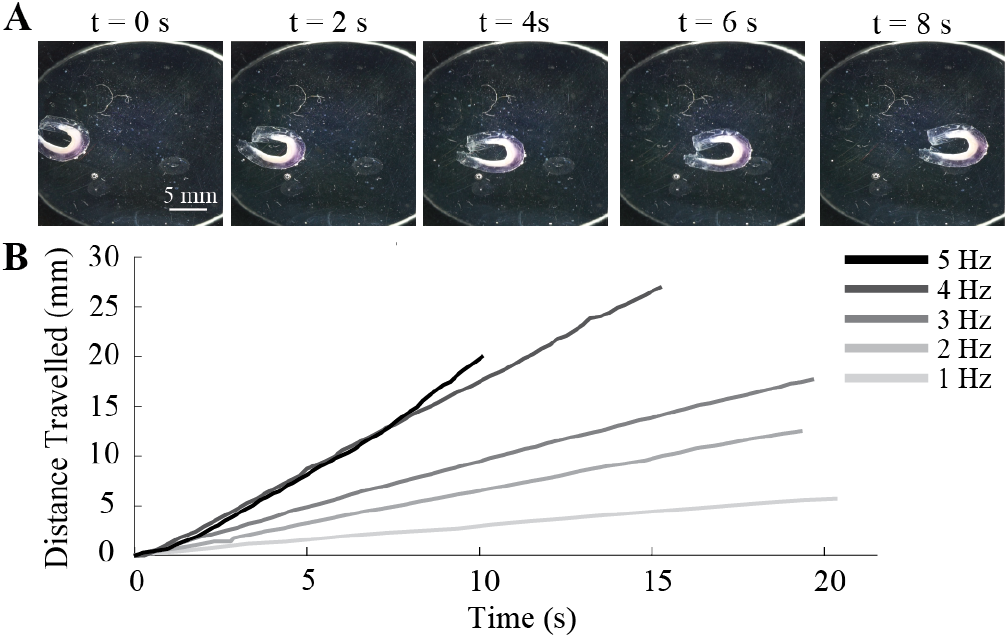
Characterization of the speed of a bio-actuator at varying stimulation frequency. (A) Time-series of bio-actuator movement at 5 Hz stimulation (3.3 V/cm, 1% duty cycle). (B) Results of tracking of bio-actuator distance traveled stroke-by-stroke over time for 5 different stimulation frequencies. Different lines correspond to different frequencies, as indicated in the legend.

Increasing the stimulation frequency increases the actuation frequency of the bio-actuator, which could account for the increase in speed. To better investigate the effect of stroke frequency on the movement of the bio-actuator, we analyzed the magnitude of the stroke by measuring the distance between the two ends of the bio-actuator (Figure 6). The closed state was taken to be 0 mm differences in all cases (Figure 6). Statistical values were obtained by averaging across 10 consecutive strokes of the bio-actuator. The stroke magnitude at 5 Hz (2.9 ±0.2 mm) is lower than the stroke magnitude of all other frequencies, including at 4 Hz. At an actuation frequency of 5 Hz, the actuated muscle does not completely recover to a resting state. Hence, increasing the actuation frequency from 4 to 5 Hz affects the speed less since the increase in stroke rate is compounded by a decrease in stroke magnitude. Statistical analysis using oneway ANOVA and Tukey’s post-hoc tests confirms that there is a statistical difference (α*<*0.05) between the 5 Hz stroke magnitudes and all other groups. We further find that there is no statistically significant difference between the stroke magnitudes of the 1 Hz and 2 Hz groups, as well as the 3 Hz and 4 Hz groups. The statistical differences observed between the 1 and 2 Hz and 3 and 4 Hz stroke magnitudes may be due to an effect of drag or inertia on the contraction movement of the bio-actuator. Further modeling of the fluid dynamics of the bio-actuator would be necessary.

**Fig. 6.**
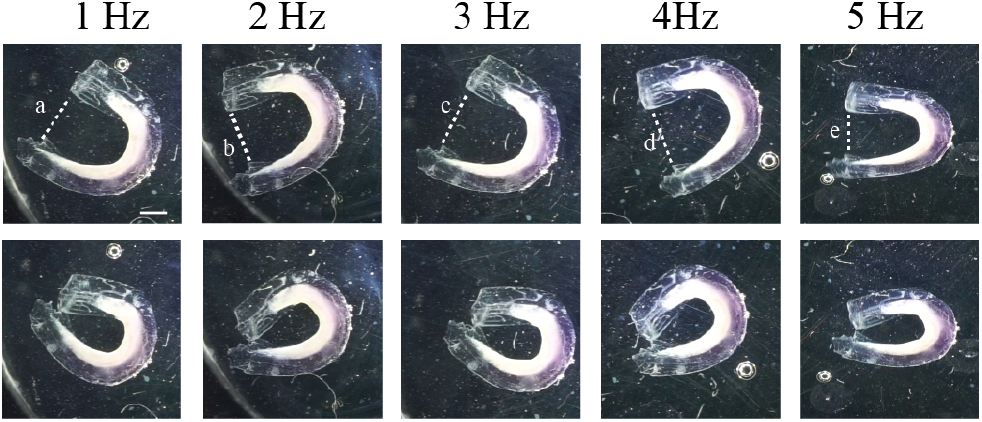
Characterization of the stroke of the bio-actuator. The top row images are the open configuration of the stroke; the bottom row images are the closed configuration. Images at each frequency are paired in columns as labeled. The dotted lines correspond to the distance that was measured. The average values were calculated based on ten consecutive strokes. (a) 3.9±0.05 mm; (b) 3.9±0.07 mm; (c) 4.5±0.1 mm; (d) 4.6±0.2 mm; (e) 2.9±0.2 mm. Scale bar is 2 mm.

**Fig. 7.**
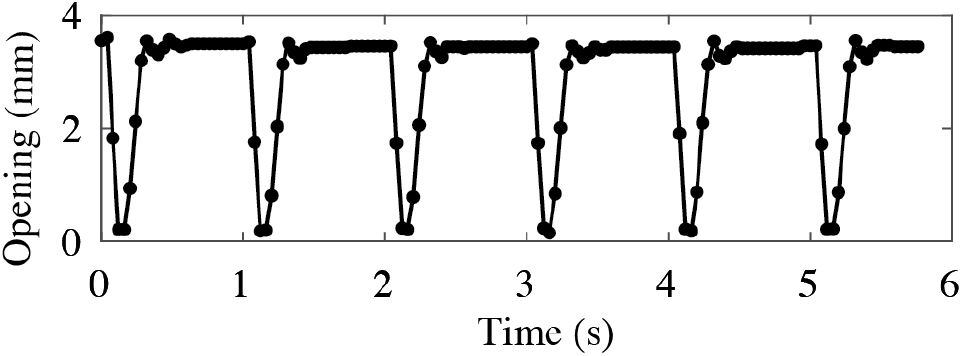
Analysis of the swimming stroke of the bio-actuator. The opening between the two ends is shown over time by analysis frame-by-frame videos for 6 consecutive strokes. Video speed is limited to 25 fps, hence each frame is taken to correspond to 40 ms.

A mechanistic understanding of the bio-actuator motion was achieved by analyzing the symmetry of the contraction and expansion. The Reynold’s number of our bio-actuators is 15, which potentially falls in the regime where intertial effects contribute to increased speed despite the reciprocal motion of the actuator. Further, we find that the contraction of the swimmer is faster than the expansion, as shown by frame-by-frame analysis of 1 Hz swimming motion. Video analysis was limited to 25 fps (40 ms per frame). (Figure 6) shows that the contraction of the bio-actuator occurs over 2 frames (approx. 80 ms), while recovery requires 3 - 4 frames (approx. 120 - 160 ms). The contraction speed is governed by an active force of the muscle, while expansion is governed by a spring-like recovery of the scaffold.

### C. Biological Characterization of the Muscle Layer

Figure 8 shows the phenotype and maturity of the myofibers of the skeletal muscle layer by immunohistochemical staining with DAPI for nuclei, F-actin to stain myofibers, and MyHC, to stain fully differentiated myofibers. The F-Actin stain in Figure 8A shows a collection of short and randomly aligned myofibers, which are not fully mature. However, the MyHC stain in Figure 8B of the same tissue segment confirms the presence of differentiated, multinucleated myofibers in the direction of groove alignment. Therefore, the bio-actuator consisted of a mix of mature and immature myofibers. It is possible that the mature myofibers were in contact with the topography of the scaffold, while the upper layer is immature cells that did not have sufficient topographical cues to form anisotropic myofibers. To confirm that anisotropy affected tissue maturation, more investigation down to the microscale analysis is needed, which could correlate the observed cell phenotypes with the matrix mesoarchitecture.

**Fig. 8.**
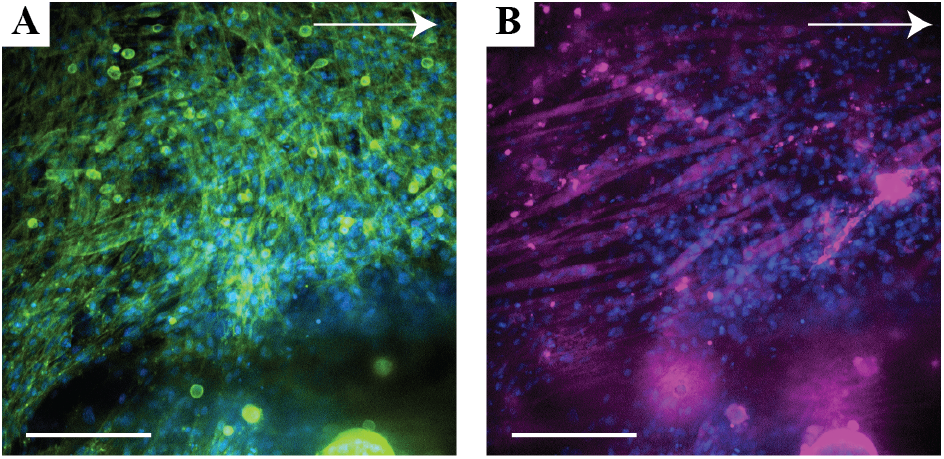
Characterization of the layered actuator (arrow indicates groove orientation). (A) F-Actin (green) and DAPI (blue) staining of myofibers. (B) MyoHC (purple) and DAPI (blue) staining of myofibers. Scale bars are 100 μm.

### D. Bio-Actuator Speed Comparison to State of the Art

Figure 9 compares the locomotion speed achieved by our skeletal muscle bio-actuator with speeds achieved by previously reported bio-actuators. Our layered planar bio-actuator achieves a speed that is at least 3.4x faster than any other bio-actuator realized from skeletal muscle tissue (Figure 9A). Our planar layered bio-actuator outperforms almost all of the cardiomyocyte swimmers, which are generally faster than skeletal muscle crawlers or walkers (Figure 9B). The only biohybrid robot reported that is faster than our bio-actuator is a fish with a pair of antagonistic cardiac muscle layers for actuation [7]. However, we are confident that with further optimization of the muscle maturation protocol and swimmer design of our skeletal bio-actuators, we can reach similar reported speeds.

**Fig. 9.**
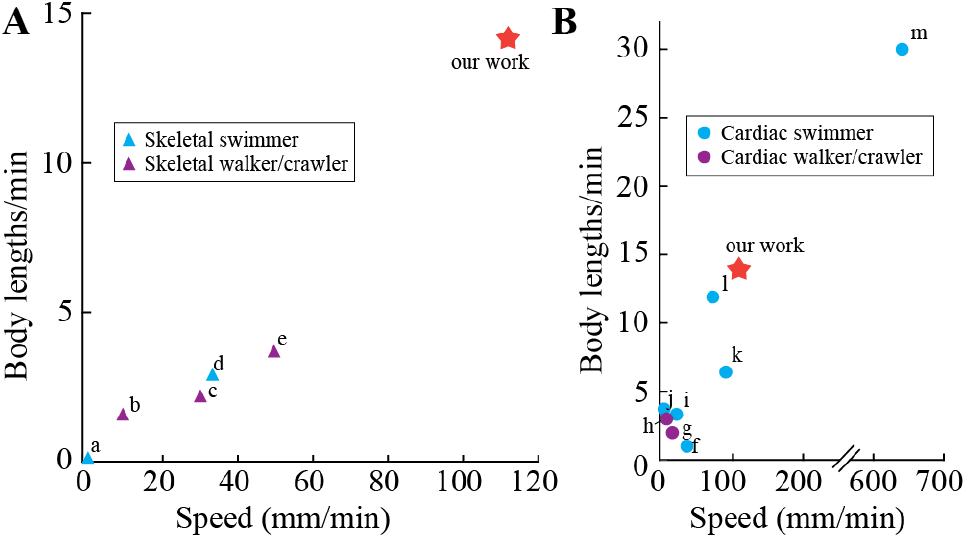
Comparison of our skeletal-muscle-based swimmer to existing biohybrid locomotion systems. (A) Reported skeletal muscle bio-actuators based on speed (mm/min) on the x-axis and body lengths/min on the y-axis. The bio-actuator from this work is shown with a star. (B) Reported cardiac muscle bio-actuators graphed based on speed (mm/min) on the x-axis and body lengths/min on the y-axis. The comparison of our skeletal muscle bio-actuator, though from different cell types, from this work is shown with a star. Literature is referenced as follows: (a) [16], (b) [9], (c) [17], (d) [10], (e) [11], (f) [18], (g) [19], (h) [20], (i) [4], (j) [21], (k) [6], (l) [5], (m) [7].

Planar geometries are particularly well-suited for locomotion through media compared to leg-based crawling systems. A layered planar geometry increases the particle’s characteristic length and opposing drag force. Most of the previously reported skeletal muscle bio-actuators have had a Reynolds number Re *<* 5, which falls in the regime where both inertial and viscous forces affect the particle motion [10]. The Re of our bio-actuator is higher, at 15, potentially falling in the regime where inertial effects contribute to increased speed. Importantly, the fastest cardiac constructs have Re = 20, 21, and 242, thus confirming that inertial effects might have a crucial role in producing high speed.

## IV. Conclusions and Future Work

In conclusion, we developed a bilayered biofabrication approach for a highly performant planar skeletal bio-actuator. We use micro-grooved hydrogel scaffolds optimized for cell adhesion and structural stability. We achieve strong interface adhesion between the cell matrix and scaffold materials, ensuring bio-actuator structural stability throughout the differentiation process.

Our bio-actuators are capable of biomimetic undulation and bending-like deformations, which, due to biofabrication constraints, have been limited to cardiac tissue-based systems. By outperforming all previously reported skeletal muscle biohybrid systems and almost all cardiac muscle tissue-based actuators, our bio-actuator demonstrates that our bilayered biofabrication approach can unlock the potential of skeletal muscle tissue for efficient locomotion within fluids.

Moreover, the finding that our constructs are composed of a mix of mature and immature myofibers further expands their potential as bio-actuators. Optimization of the maturation process by varying the volume of the cell matrix and the topographical pattern will allow better maturation of myofibers. Further, we show results from one optimal bio-actuator; optimization of the maturation process will also ensure reproducibility. Hence, our bio-actuators could become even more powerful and performant, and potentially enable novel bio-actuator design modularity to realize robots that combine multiple muscle tissue building blocks for yet unexplored complex motion.

## ACKNOWLEDGMENT

The authors acknowledge Niklas König (xolo3D) for contributions with xolographically-printed hydrogel formulation. This work was done within the framework of the ALIVE initiative (Advanced Engineering with Living Materials) and funded by the SFA-AM program (Strategic Focus Area – Advanced Manufacturing).

